# PA200 differentially regulates the proteasome and inhibits migration of cancer cells

**DOI:** 10.1101/2024.12.17.628836

**Authors:** Ayse Seda Yazgili, Georgia A. Giotopoulou, Sabine J. Behrend, Frauke Koops, Vanessa Welk, Thomas Meul, Linda Zemke, Norbert Reiling, Torsten Goldmann, Georgios T. Stathopoulos, Silke Meiners

## Abstract

Proteasome activator 200 (PA200) is upregulated in non-small cell lung cancer (NSCLC) and linked to poor prognosis. We previously demonstrated that the overexpression of PA200 in NSCLC is associated with immune evasion and reduced responsiveness to immune checkpoint inhibitors. The cell autonomous function of PA200 in cancer growth, however, is not solved.

We here demonstrate that deletion of PA200 in two distinct lung cancer cell lines induced cell-specific alterations in proteasome composition and activities with a minor direct impact on overall proteasome activity. Deficiency of PA200 in lung cancer cells did not consistently alter tumor cell growth *in vitro* and *in vivo*. However, we observed concerted inhibition of tumor cell migration and invasion with conserved downregulation of the integrin ITGB3 and transcriptional dysregulation of multiple cell adhesion and ECM regulators. Our transcriptome profiling revealed a striking disparity in the transcriptional response to PA200 deletion in the two lung cancer cell lines. Together with our PA200 interactome analysis that uncovered an unexpected cell-dependent profile of PA200-interacting proteins, our data indicate that the function of PA200 is cell-specific and depends on the cellular context. In conclusion, we here demonstrate that PA200 cell-autonomously regulates the invasive capacities of tumor cells thereby potentially promoting lung cancer spread and metastasis formation. This mechanism might add to PA200-related immune evasion and may contribute to the observed poor prognosis of PA200-overexpressing lung cancer patients.

## INTRODUCTION

The proteasome system is key for regulated intracellular protein degradation. It consists of multiple proteasome complexes that share a common proteolytic core, the 20S proteasome, but differ in their associated proteasome activators which fine-tune substrate degradation by the proteasome ^1^. The 20S proteasome contains three active sites that cleave the proteins after hydrophobic, basic, and acidic amino acids constituting the chymotrypsin-like, trypsin-like and caspase-like proteolytic activities, respectively ^2^. These active sites can be exchanged by inducible subunits, i.e. LMP7, MECL1, and LMP2 to assemble into the immunoproteasome that has altered proteolytic activities ^3^. The proteasome activators modulate proteasome function in specific cellular compartments and in an ATP- and ubiquitin-dependent or -independent manner ^1^. The well-known 19S regulator binds to the 20S proteolytic core either on one or both ends forming the 26S and 30S proteasome, respectively. These proteasome complexes mediate ATP- and ubiquitin-dependent protein degradation ^4^. Yet, alternative proteasome activators including PA28αβ, PA28γ, and PA200 act ATP- and ubiquitin-independently and have distinct subcellular functions ^2^.

PA200, encoded by the PSME4 gene, is expressed at low levels in most cells of the organism except for germ line cells ^5^. PA200 has been implicated in various cellular processes, such as DNA damage repair ^6,7^, mitochondrial and proteostasis stress responses ^8,9^, myofibroblast differentiation ^10^, aging ^11,12^, and protection from neuropathy ^13,14^. It is upregulated in lung fibrosis, melanoma and lung cancer ^10,15,16^. PA200 was recently shown to restrict antigenic diversity of non-small cell lung cancers (NSCLC) favoring immune evasion and responsiveness to immune checkpoint inhibition ^15^. The cell-autonomous role of PA200 in cancer, however, remains elusive.

In this study, we investigated the functional role of PA200 in lung cancer by genetically deleting PA200. We used two distinct lung cancer cell lines, A549 and H1299, and characterized them thoroughly regarding proteasome regulation, cellular growth and migration capacity. Our findings revealed a striking disparity of PA200 deficiency on proteasome complex regulation and transcriptional profile for the two cancer cell lines, however cancer cell growth was not consistently altered by PA200 knockout. In contrast, we found that PA200 deletion resulted in decreased cell migration and extracellular matrix (ECM) interactions consistently in both cell lines. Our data suggests that PA200 plays a cell-autonomous role in the invasive capabilities of tumor cells, which may contribute to the spread of lung cancer and the formation of metastases. This mechanism might add to PA200-related immune evasion and may contribute to the observed poor prognosis of PA200-overexpressing lung cancer patients ^15^.

## RESULTS

### Deletion of PA200 in lung cancer cell lines differentially impacts proteasome function and assembly

We first confirmed the oncogenic role of PA200 in lung cancer by analysis of PA200 protein expression in different lung cancer types, i.e., lung adenocarcinoma (LUAD), squamous cell carcinoma (SQCLC), and small cell lung cancer (SCLC), in a cohort of 60 patients using immunohistochemistry with a validated antibody ^10,17^. Expression of PA200 is generally low in lung parenchymal and immune cells, as previously demonstrated by us ^10^. We observed significantly increased expression of PA200 in LUAD and SQCLC compared to SCLC tumors (Suppl. Figures S1A-B). PA200 was also strongly upregulated in a genetic model of Kras-induced lung adenocarcinoma in mice (Suppl. Figure S1C) supporting its induction upon oncogenic transformation of epithelial cells ^18^.

To elucidate the functional role of PA200 in cancer, we deleted PA200 in two lung cancer cell lines, i.e. A549 and NCI-H1299 (hereinafter H1299) using CRISPR/Cas9 technology. Complete loss of PA200 was confirmed for several A549 and H1299 cell clones by RT-qPCR (Figure 1A), Western blotting (Figures 1B-C) and DNA sequencing (Suppl. Figure S1D). We noted that H1299 cells express more PA200 than A549 cells, as observed for mRNA (Figure 1A) and protein levels (Figure 1B). Unexpectedly, deletion of PA200 in the two cell lines affected proteasome activity differentially: In A549 cells, both the total chymotrypsin-like (CT-L) and total caspase-like (C-L) activities were significantly reduced compared to WT controls (Figure 1D), while H1299 PA200 knockout (KO) cells displayed significant activation of these two activities (Figure 1E). Contrary to previous findings ^15,19^, loss of PA200 did not affect the total trypsin-like (T-L) activity (Figures 1D-E). To analyze these cell-specific effects of PA200 on the proteasome in more detail, we performed native gel analysis to determine the activity, composition, and abundance of distinct proteasome complexes ^20^. In A549 PA200 KO cells, the CT-L activity of the 20S was slightly activated while the activity of 26S and 30S complexes were not significantly altered (Figure 2A). In H1299 PA200-deficient clones, 20S activity was reduced while 30S activity was elevated (Figure 2B). Blotting of the native gel and probing for the α-20S subunits (α1-7), revealed no significant changes in the abundance of the proteasome complexes upon PA200 deletion in A549 cells (Figure 2A) but elevated abundance of 30S proteasome complexes in H1299 PA200 knockout clones accounting for their elevated 30S activity (Figure 2B). Of note, we observed a strikingly divergent composition of proteasome complexes upon deletion of PA200 (Figures 2A-B). PA200-deleted A549 clones assembled higher levels of immunoproteasome subunits in the 20S (for LMP7) and 26S and 30S complexes (for MECL1 and LMP2) and had higher levels of PA28 activator bound to the 20S (Figure 2A). The increased assembly of the immunoproteasome could explain the reduced C-L activity in PA200-deleted A549 clones (Figure 1C). In contrast, incorporation of the immunoproteasome subunits was significantly decreased in PA200-deleted H1299 cells (Figure 2B). The levels of PA28α and PA28β were not significantly regulated in H1299 (Figure 2B), however PA28β was upregulated significantly in A549 PA200 KO cells (Figure 2A). Our results suggest that the deletion of PA200 has a minor direct impact on overall proteasome activity but rather contributes to cell-specific adjustment of proteasome composition and activities.

**Figure 1:**
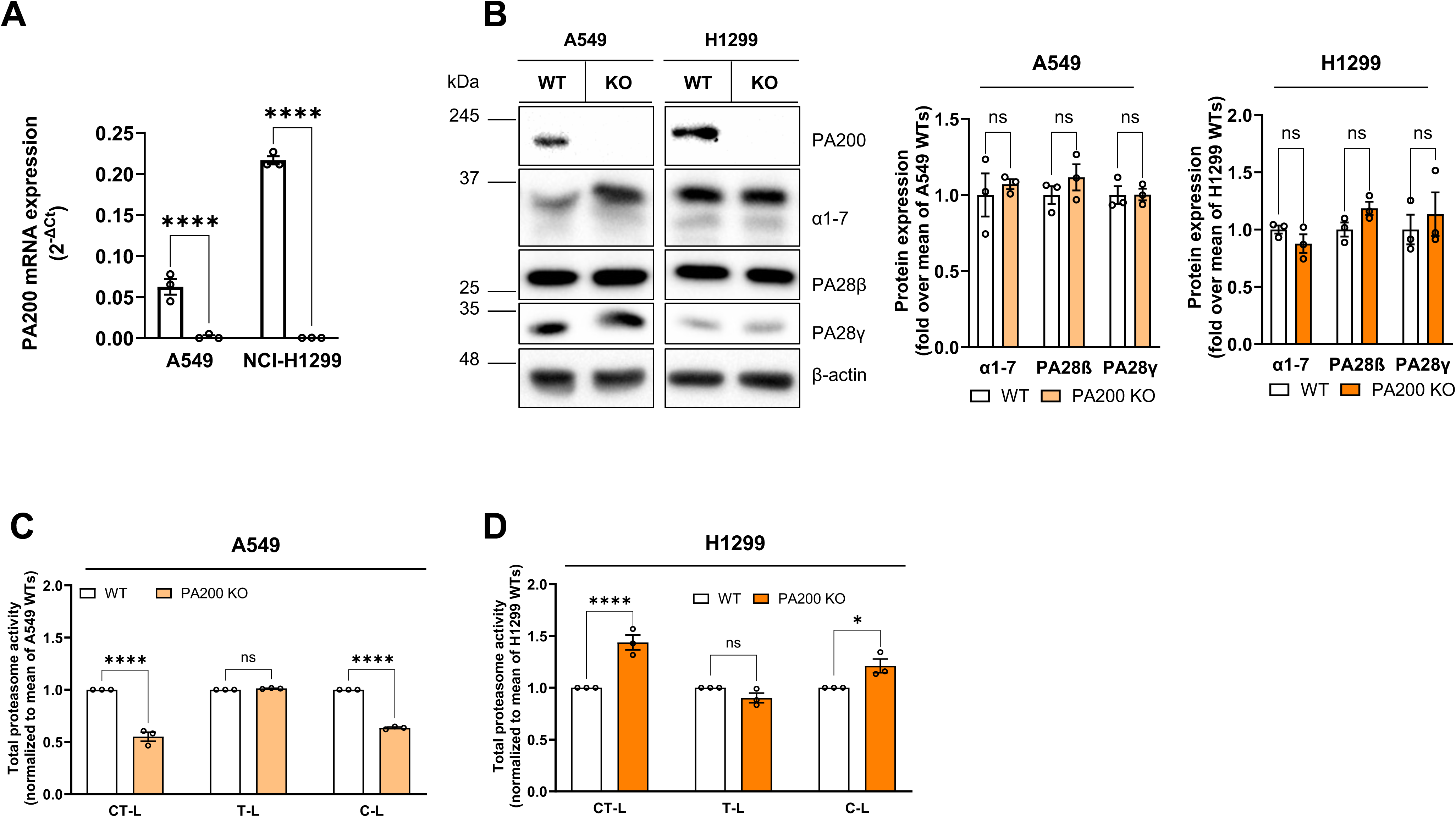
Deletion of PA200 in lung cell lines differentially impacts proteasome activity. PA200 KO was generated by CRISPR/Cas9 in A549 and H1299 cell lines. A) Relative PA200 mRNA expression was determined in A549 and H1299 WT and KO cell lines by RTqPCR using RPL19 as a housekeeping mRNA (2^-deltaCT^ method, ****p < 0.0001, ordinary 2-way ANOVA for analysis of WT vs KO and unpaired t test for A549 and H1299 comparison, n=3 clones). B) Representative images and relative protein expression levels as determined by Western blotting. β-actin served as a housekeeping protein to ensure equal protein loading. Total proteasome activity of CT-L, T-L and C-L was analyzed in the different clones of C) A549 and D) H1299 (normalized to the respective WT control clones).

**Figure 2:**
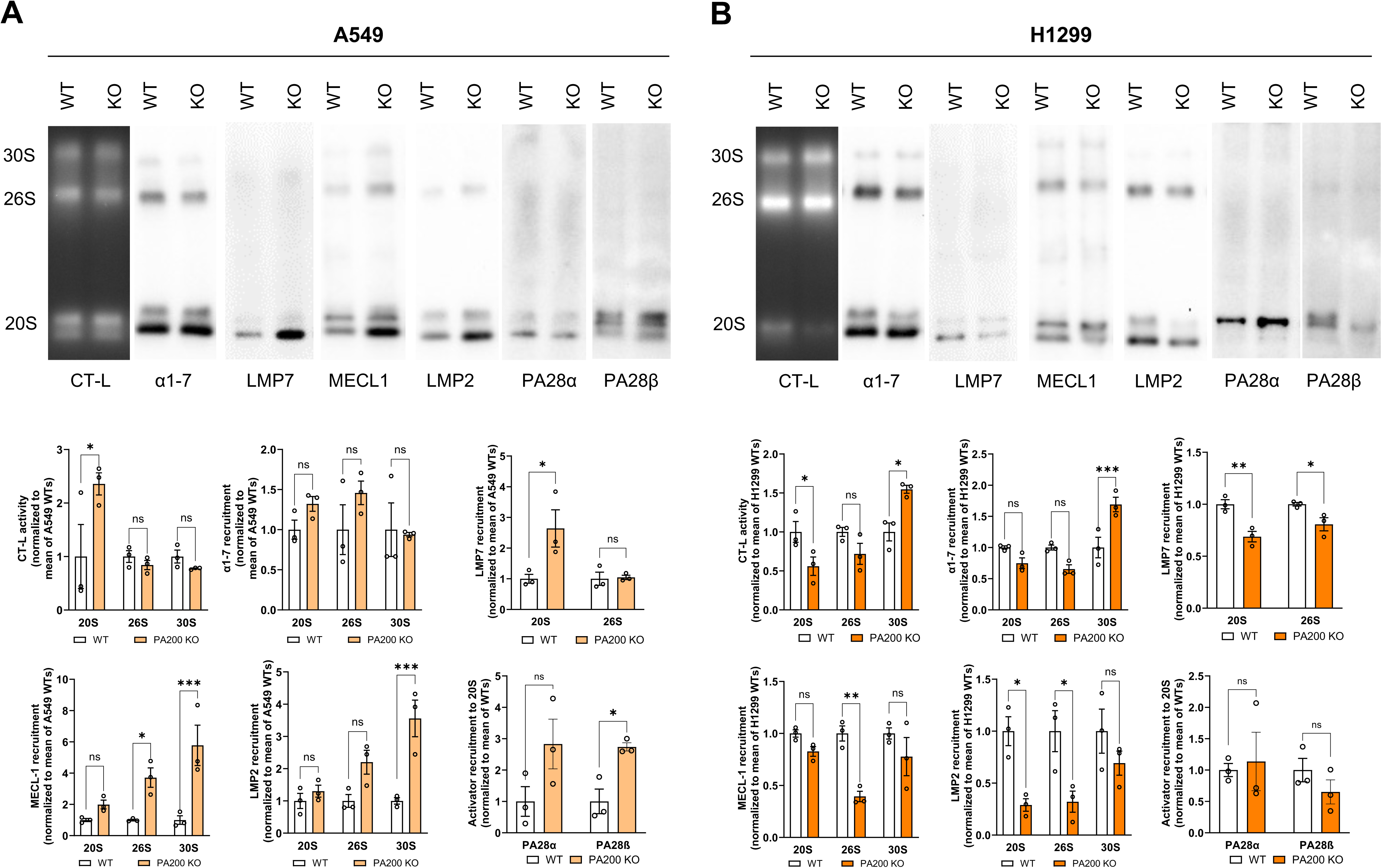
PA200 KO differentially affects proteasome composition and activity in lung cancer cell lines. The composition and activity of proteasome complexes was compared between the selected WTs and KO cell lines. The gels were blotted after the in-gel proteasome CT-L activity of native gels to determine the distribution of catalytic subunits and activators subunits in proteasome complexes of (A) A549 cells and (B) NCI-H1299 cells. Quantifications are shown below the respective images. (*p < 0.05, **p < 0.01, ***p < 0.001, ordinary two-way ANOVA with Šídák test, n=3, different passages).

### PA200 deficiency does not alter tumor cell growth in vitro and in vivo

We next aimed to investigate the cellular consequences of PA200 deletion and differential proteasome regulation and investigated the growth of the lung cancer cell lines. In A549 cells, deletion of PA200 slightly reduced proliferation rates (Figure 3A) but did not affect colony formation (Figure 3B). H1299 cells did not show any significant alteration neither in proliferation (Figure 3A) nor in colony formation assays (Figure 3B). To analyze tumor cell growth *in vivo*, we injected pools of A549 and H1299 PA200 WT and KO cell clones into the flanks of immunocompromised SCID mice and monitored tumor growth and body weight for 8 weeks. SCID mice are deficient in B and T cells and therefore lack adaptive immune responses. This model thus allows us to dissect the cell-autonomous function of PA200 in tumor growth in the absence of its immune-modulating function ^15^. The development of primary tumors was reduced for A549 PA200 KO cells (Suppl. Figure S2A) while all animals with H1299 WT and PA200 KO cells injected developed tumors. For both cell lines, we observed delayed flank tumor development of PA200 deleted cells. Tumor growth rates, however, normalized over 60 days and were not significantly different from respective WT controls at the endpoint of analysis (Figures 3C-D, Suppl. Figures S2B-E). Ki-67 staining of primary tumors did not reveal any significant difference in the number of proliferating cells between the WT and PA200-deleted tumors (Figures 3E-F). These *in vitro* and *in vivo* data demonstrate that PA200 does not consistently alter tumor cell growth in a cell autonomous manner.

**Figure 3:**
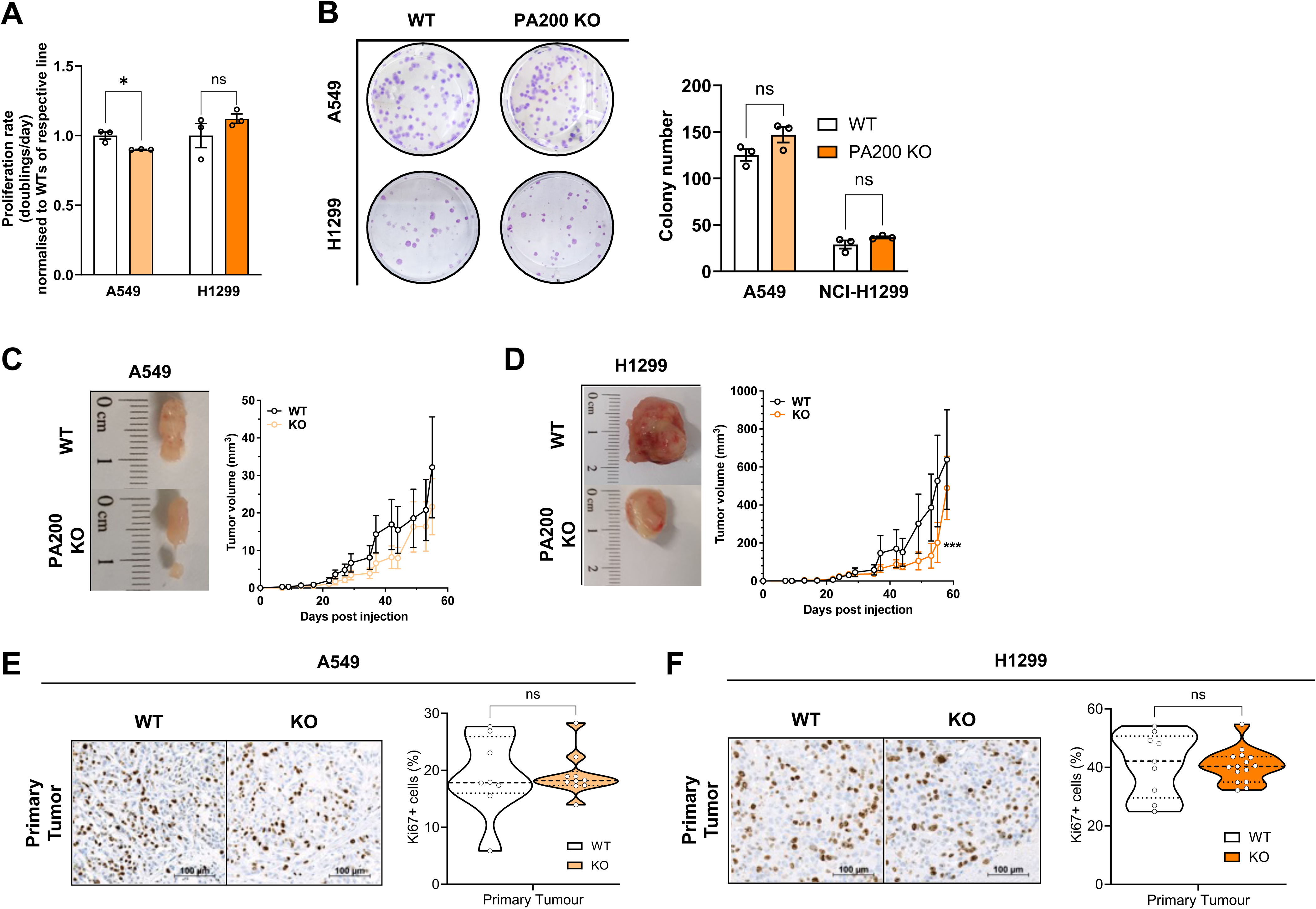
Deletion of PA200 in lung cancer cell lines does not consistently alter tumor cell growth. A) Pools of three A549 and H1299 WT and KO cell clones were seeded and counted on different days to track the doubling time. Data were normalized to the respective WT control (n=3, different passages, unpaired t-test). B) A clonogenic assay optimized for A549 and H1299 pools of clones. Visible colonies that had at least 50 cells were counted and quantified (n=3, different passages, unpaired t-test). Primary tumor volumes of C) A549 and D) H1299 cells were monitored over time (ordinary 2-way ANOVA, WT cells injected into 10 mice and PA200 KO cells injected into 15 mice, *p < 0.05, ***p < 0.001). Ki67 staining of dissected primary tumors from E) A549 and F) H1299 WT and respective KO cells (unpaired t-test).

### PA200 deficiency inhibits tumor cell migration and invasion capacity in vitro

We further characterized the effects of PA200 deletion with regard to another hallmark of cancers, i.e. their capacity to migrate and invade the extracellular matrix (ECM) ^21^ and analyzed the effect of PA200 deletion in wound healing and migration assays. Loss of PA200 in A549 cells reduced invasion of matrigel-embedded A549 spheroids (Figure 4A). As H1299 cells are unable to form spheroids, we assessed their migration efficiency in a Boyden chamber assay and showed prominent loss of migration capacity upon PA200 deletion (Figure 4B). Using a standardized wound-healing assay, we confirmed reduced migration and wound closure of both A549 and H1299 PA200 KO cells (Figures 4C-D). We next assessed whether deletion of PA200 promotes epithelial-mesenchymal transition (EMT), a hallmark feature of invasive epithelial tumor cells ^21^. Upon KO of PA200, the epithelial adherens junction protein E-cadherin but also the mesenchymal N-Cadherin were significantly upregulated on the protein level in both A549 and H1299 cells (Figure 4E). Expression of the tight junction component claudin and the mesenchymal markers, such as vimentin and β-catenin, was not regulated by PA200 deficiency (Figure 4E). Deletion of PA200, however, induced strong and concerted downregulation of Integrin-β3 (ITGB3) expression on RNA and protein levels in both lung cancer cell lines (Figure 4E and Suppl. Figures S3A-B). Taken together, our results indicate that PA200 plays a conserved role in the regulation of cell migration and invasion of lung cancer cells, potentially related to downregulation of ITGB3.

**Figure 4:**
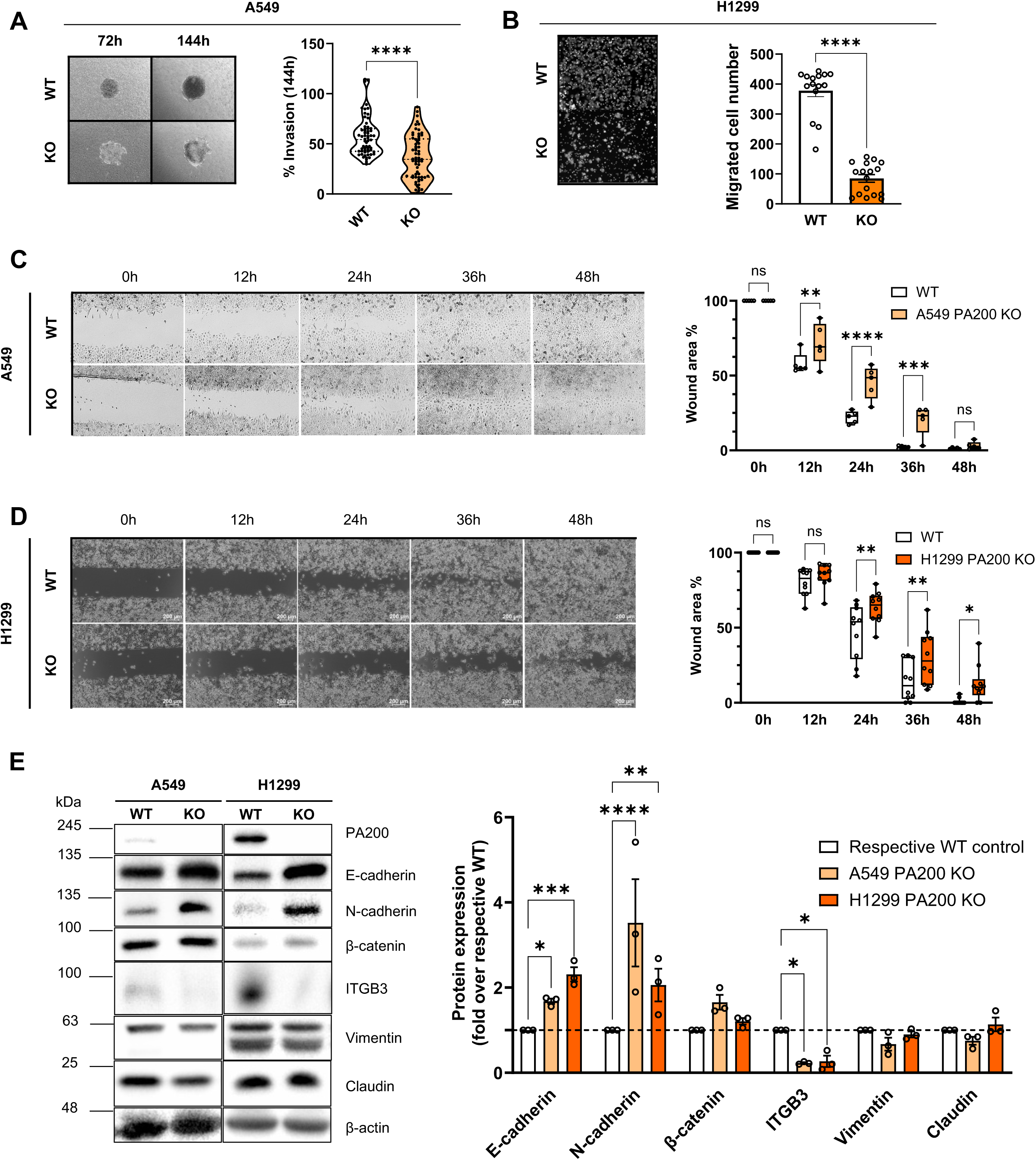
Deletion of PA200 regulates invasion and migration capacity of lung cancer cells. A) An 3D invasion assay was employed for A549 WT and KO cell pools to determine their invasive capacity. Representative images of the spheroids after 72 h of initial seeding and 72 h after embedding in the Collagen G+Matrigel membrane (left panel). The invasion percentage was calculated by comparing the 72h and 144 h area differences (right panel, n=3, different passages). B) Representative images of the Boyden chamber migration assay of the H1299 cells (left panel) and quantifications (right panel) are shown (n=3, different passage, two-tailed unpaired t-test). Representative photos of the wound closures of (C) A549 and (D) H1299 WT and KO cells were taken every 12 h up to 48 h and quantified (right panel, ordinary 2way ANOVA, n=3, different passages). E) Protein levels of epithelial and mesenchymal markers were quantified (right panel, n=3, different passages, 2-way ANOVA) in A549 and H1299 WT and KO pools and β-actin (representative image) served as protein loading control. For quantification, relative expression levels were normalized to the respective WT controls of each cell line. *p < 0.05, **p < 0.01, ***p < 0.001, ****p < 0.0001.

### Transcriptomic analysis highlights conserved downregulation of ECM-related genes in PA200-deficient lung cancer cells

To disentangle cell-specific from conserved effects of PA200 deletion, we performed transcriptional profiling of our WT and PA200 KO A549 and H1299 cell clones. The transcriptomes of WT and KO clones were clearly distinct as highlighted by PCA analysis (Suppl. Figures S4A-B). Deletion of PA200 had a more substantial transcriptional effect in A549 compared to H1299 cells with 345 genes being down- and 419 up-regulated in PA200-deleted A549 cells and 288 genes down- and 196 genes up-regulated in H1299 PA200 KO clones (Figures 5A-C). Surprisingly, we found only 16 genes that were co-regulated both in PA200-deleted A549 and H1299 clones (highlighted in red (up) and blue (down)). These included, besides PSME4, several proteins involved in cytoskeletal organization, cell adhesion and remodeling, and interaction with ECM. Amongst them, we confirmed ITGB3 as concertedly downregulated in both lung cancer cell lines (Figure 5C). Reactome pathway and molecular signature database (MsigDB) analysis of differentially regulated genes confirmed that PA200 deletion in both A549 and H1299 cell lines, concertedly regulated pathways related to ECM formation such as collagen formation, ECM organization, proteoglycans, and integrin and surface organization-related genes (Figures 5D-G, Suppl. Figure 4C-D). Our transcriptional analysis thus confirms a conserved role for PA200 in the regulation of invasion and migration that is mediated by transcriptional regulation of cell adhesion and ECM regulators. At the same time, the minimal overlap of the transcriptional response to PA200 deletion in the two NSCLC cell lines indicates that the function of PA200 is cell-specific and depends on the cellular context.

**Figure 5:**
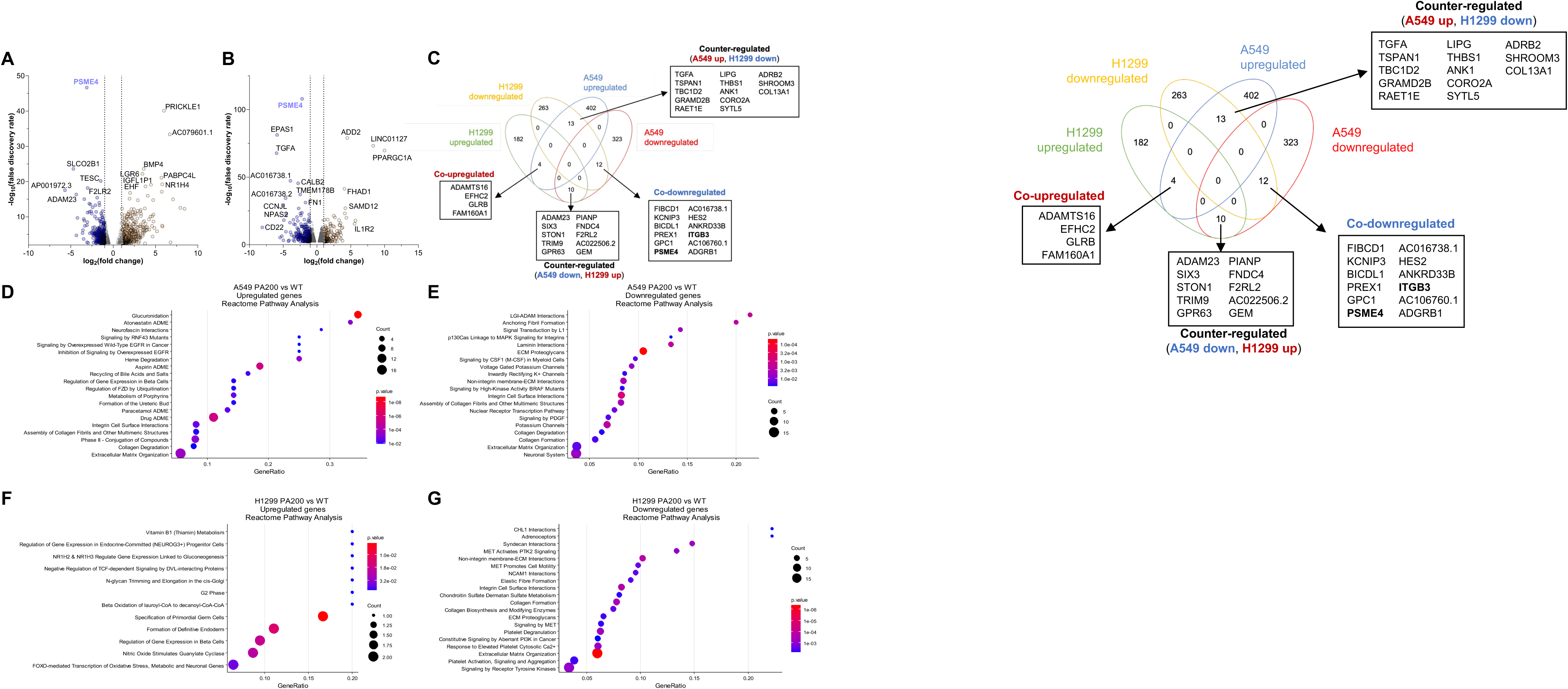
RNA sequencing analysis reveals differential transcriptome regulation upon PA200 deletion in lung cancer cell lines. The effects of PA200 deficiency in A549 and H1299 on transcriptional regulation were analyzed by RNA sequencing. The results were filtered, and differentially expressed genes (DEGs) (|FC| ≥ 1, p ≤ 0.05, FDR ≤ 0.05) were plotted in volcano plots for A) A549 and B) H1299 cell lines. C) Venn diagram of the co-regulated and counter-regulated DEGs showed transcriptomic differences between the two lines. Bubble charts of Reactome analysis showed the significantly up (left panel) or down (right panel) regulated pathways in D-E) A549 and F-G) H1299 cells. (n=3, WT and KO).

### Pulldown of PA200 reveals differential interactome in lung cancer cell lines

These prominent cell-specific effects of PA200 deletion led us to assume that PA200 has differential roles in the two cancer cell lines that might be revealed by its interactome. To determine PA200 interactors, we immunoprecipitated PA200 from A549 and H1299 WT cell lines. Interacting proteins were identified by mass-spectrometry. Immunoprecipitation of PA200 from the respective PA200 KO clones was used as a specificity control (Figure 6A). Our interactome analysis of PA200 identified 91 (A549) and 41 (H1299) interacting proteins with only a minimal overlap of three proteins including PA200 (PSME4) (Figure 6B). We were surprised not to detect any significant enrichment of proteasomal subunits in our mass-spectrometry data, even though 20S subunits were clearly enriched in the PA200 pulldowns (Figure 6A). This might be due to the weak unspecific binding of the proteasome to the anti-PA200 antibody (Figure 6A) which could explain why proteasome subunits were not detected as significantly enriched when we compared interacting proteins between WT and respective KO clones. In the next step, we analyzed our interacting proteins with MSigDB and Reactome databases to understand the functions and pathways of the PA200 interactors. In A549 cells, the interactors of PA200 were mainly related to E2F and Myc targets, reactive oxygen species, ultraviolet (UV) response, glycolysis, and G2-M checkpoint proteins (Figures 6C and 6E). Reactome pathway analysis showed that the PA200 interactors in A549 cells were mainly involved in DNA processes (Suppl. Figure S5A). PA200 interacting proteins in H1299 cells were related to the molecular signatures of the mitotic spindle, UV response, mTORC1 and PI3K/AKT/mTOR signaling (Figures 6D and 6F). In the Reactome analysis, they showed enrichment in Rho GTPase signaling and regulation of transcription (Suppl. Figure S5B). Concludingly, these data reveal an unexpected cell-specific profile of PA200 interacting proteins which might explain the differential cellular response to PA200 deletion in A549 and H1299 lung cancer cell lines.

**Figure 6:**
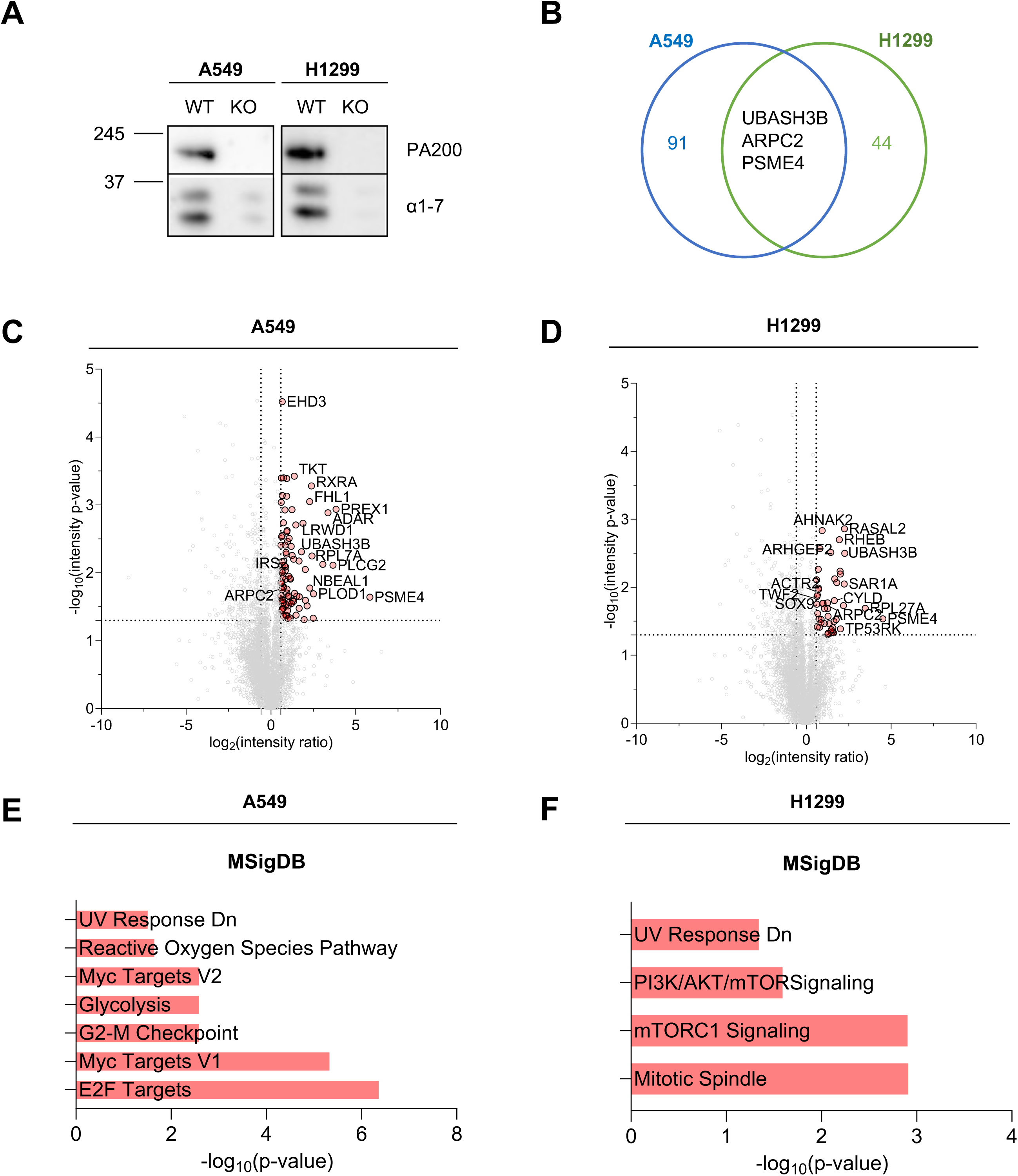
Differential interactome of PA200 in lung cancer cell lines. Interactors of PA200 were captured by PA200 immunoprecipitation and analyzed by mass spectrometry. PA200 KO cell lines were used as control lysates. A) Representative images of the Western blot analysis of the PA200 immunoprecipitation with A549 and H1299 cell lysates, with α1-7 and PA200 antibodies. B) Venn diagram showing the significantly enriched proteins in pulldowns of PA200 (PSME4) in A549 and H1299 cell lines. Volcanoplot showing all significantly enriched interacting proteins of PA200 compared to control lysates for C) A549 and D) H1299 cells in light pink circles (FC ≥ 1.5, p ≤ 0.05). The PA200 interactors were analyzed with EnrichR to highlight the molecular signature (MsigDB) differences of E) A549 and F) H1299 PA200 interactomes. LC-MS/MS was performed by Dr J. Merl-Pham, Research Unit Protein Science and Metabolomics and Proteomics Core Facility (HMGU, n=3 different passages).

## DISCUSSION

We here uncover adaptive cellular responses to PA200 deletion that depend on the cellular context and are cell line specific. In our comprehensive analysis of two different lung cancer cell lines, we discovered an almost entirely distinct interactome of PA200 which coincided with differential cellular and transcriptional responses to genetic deletion of PA200. PA200 deficiency had cell-specific effects on the proteasome complex system resulting in distinct alterations in proteasome complex composition and activity. Among the conserved responses to PA200 deletion in the two cancer cell lines, we discovered cell-autonomous downregulation of cell migration and invasion upon transcriptional alterations in cell-adhesion and ECM-related genes such as ITGB3.

Deleting PA200 altered the activity and composition of proteasome complexes but with opposite outcomes depending on the cell line: While total CT-L and C-L activities were decreased in A549 cells, they were increased in H1299 cells upon PA200 deletion. Furthermore, in-gel proteasome activity assays revealed a shift in proteasome complex activities and composition including differential assembly of immunoproteasomes and association of PA28 regulators. These data reveal cell-specific adaptive remodeling of the proteasome system in response to PA200 deficiency. Our data are in line with previous findings from our own lab on the adaptation of the proteasome system upon transient knockdown of PA200 in primary human lung fibroblasts ^10^. Similarly, VerPlank et al. recently demonstrated that proteasomal degradation was activated in PA200 KO animals, contributing to protection from proteotoxic neuropathy ^13^. The differences between the two cell lines were not limited to proteasome activities but proteasome remodeling was associated with a differential transcriptional response. For example, inflammatory signaling was a prominent feature of PA200 KO in A549 cells, while this pathway was downregulated in H1299 PA200 KO cells. This discrepancy might explain the increased assembly of the immunoproteasome in A549 KO cells. Moreover, we noted that the interactome of PA200 between the two cell lines is very different with only two overlapping proteins. This might relate to weak and transient interactions that were stabilized upon pulldown of PA200 using a mild and non-stringent immunoprecipitation protocol or due to different subcellular localization and functional annotation of PA200-containing proteasome complexes. The latter is supported by the differential interaction of PA200 with the DNA and replication machinery in A549 cells versus transcription- and GTPase-related interactions in H1299 cells. One could also speculate that the divergent cell-type specific effects relate to the role of PA200 in DNA repair and histone degradation ^6,7,22,23^. This observation aligns with the finding that inositol-6-phosphate (IP6), a nuclear regulator of enzyme function and DNA repair ^24^, is an integral component of PA200 as evidenced by structural analyses (Toste Rêgo and da Fonseca 2019; Guan et al. 2020). Furthermore, the mutational background (A549 with KRAS activating mutation, H1299 loss of TP53), the origin (A549 primary lung adenocarcinoma ^26^ and H1299 metastatic lymph node ^27^ and the different baseline PA200 expression levels of the two cell lines could explain the observed functional differences. Our data thus indicate that the effects of PA200 need to be carefully analyzed depending on the cellular background.

In the lung cancer cells, we did not observe any consistent effects of PA200 deficiency on tumor cell growth, neither *in vitro* nor *in vivo*. These data are supported by our transcriptome analysis where we did not find any prominent regulation of growth related pathways upon deletion of PA200 in A549 and H1299 cells. Our data also accord with observations from PA200 KO mice that did not form tumors even when crossed to p53 mutant mice ^5^. While transient knockdown of PA200 in Hela cells increased cell growth in the presence of glutamine in a previous study ^28^, however PA200-deleted A549 and H1299 clones showed different glutamine-sensitivity (data not shown). While cell growth was unaffected, PA200 deficiency impaired migration and invasiveness of lung cancer cells. This effect was conserved in both lung cancer cell lines and corroborated using several complementary functional assays such as wound healing, Boyden chamber and Matrigel invasion assays. This decrease in the invasive capacity in PA200 deficient cancer cells can be explained by transcriptomic regulation of several cell-adhesion and ECM regulators. Although, there were very few genes that were concertedly regulated by PA200 deletion in A549 and H1299 cells, the overall transcriptomic response to PA200 deletion was similar with integrin cell surface interactions and non-integrin membrane interactions being downregulated. A prominent and concertedly reduced cell-matrix adhesion molecule was integrin beta 3 (ITGB3), a well-established driver of metastasis in several solid cancer types and associates with poor prognosis ^29–32^. We confirmed its downregulation by qRT-PCR and Western blot analysis. However, aAs our MS-based PA200 interactome analysis did not show ITGB3 as a direct PA200 interactor, dysregulation of ITGB3 might be indirect.

Taken together, our data, together with the results of the Merbl lab ^15^, point towards a tumor-promoting role of PA200 in lung cancer related to its ability to regulate cell migration, invasion, and immunogenicity.

## MATERIALS AND METHODS

### Cell culture and generation of knockout cell lines

The human lung cell lines were obtained from DSMZ or were a gift from Dr. Georgios Stathopoulos. The cell lines were cultured at 37°C and 5% CO_2_ in a humidified incubator. A549 cells (RRID:CVCL_0023) were grown in DMEM/F12 medium, while NCI-H1299 cells (RRID:CVCL_0060) were cultured in RPMI-1640 medium, both supplemented with 10% FBS and 100 U/mL penicillin/streptomycin.

Low passage number cells were harvested at 80% confluency. Transfection of TrueCut Cas9 Protein v2, Hs.Cas9.PSME4.1.AB and tracRNA (IDT, Leuwen, Belgium) was performed using the Lipofectamine CRISPRMAX Transfection Reagent protocol (Thermo Fisher Scientific). The cells were incubated at 37°C for 48 h after transfection. Following the incubation period, the cells were collected, and 50 cells were seeded onto 15 cm dishes. After 2 weeks, colonies were picked with trypsin-incubated cloning disks and transferred to 24-well (one colony per well). To confirm the knockout efficiency, protein and DNA were isolated. For PCR-based amplification we used forward primer 5’ATCTGCAAGAGAGATGCAGCC 3’ and reverse primer 5’AGGTTTGAGCAGCAGCAAGA 3’. For Sanger sequencing reverse primer 5’TGGATGGGGAAAGCAAAACCC 3’ was used.

### Cell proliferation and colony formation assays

The proliferation rate of different cell types was assessed using established methodology ^33,34^. A549 (30 000 cells/well) and NCI-H1299 (40 000 cells/well) cells were seeded into 6-well plates. The initial cell count was determined on day 1, and the final cell count was recorded on day 4 to calculate the rate of cell division per day according to the proliferation Rate formula = log₂ [(Final day cell count (Day 4) / Initial day cell count (Day 1))] / (Final day - Initial day (3 days)). A colony formation assay was conducted by seeding 100 NCI-H1299 and 200 A549 cells in 6-well plates as previously described ^35^. The culture medium was refreshed on day 3 and day 7. On day 10, the cells were fixed using 4% formaldehyde, stained with 1% crystal violet dye (Sigma), photographed, and quantified.

### Migration and invasion assays

For Boyden chamber migration analysis, a 10% FBS medium was used as a chemoattractant in 24-well plates. Boyden chambers (Biotrend, Cologne, Germany) containing 100 000 NCI-H1299 cells in 0% FBS medium were inserted into the wells. After 24 hours, the cells were fixed with 4% formaldehyde and stained with 1% crystal violet solution. Excess dye was washed off, and the chambers were left to dry before imaging with Axio Observer.Z1/7 (Zeiss, Oberkochen, Germany).

For wound healing assays, inserts (Ibidi, Graefelfing, Germany) were used to assess cellular wound healing capacity. The inserts were placed into a 24-well plate (1 insert/well), with A549 (28 000 cells/insert side) and NCI-H1299 (20 000 cells/insert side) cells seeded on both sides. Inserts were removed after 24 h and 1 ml of medium was added. For experiments related to Figure 4, H1299 cells were incubated in a live cell imaging microscope (Axio Observer.Z1/7, Zeiss) at 37°C and 5% CO_2_ for 48 h or until the 500 μm gap closed. Images from different time points were calculated using the Wound Healing Tool macro on ImageJ (RRID:SCR_003070) ^36^. A549 cells were imaged in a live cell imaging multimode reader (Cytation1, Agilent Technologies, Waldbronn, Germany) at 37°C and 5% CO_2_ for 48 h. Using a 4x objective, one to two images per well were captured at different time points using bright field imaging channels. If two pictures were taken, the images were stitched prior to analysis. Images were analyzed using the Gen5 3.16 Software (Agilent Technologies) following the application note “Incorporation of a Novel, Automated Scratch Tool and Kinetic Label-Free Imaging to Perform Wound Healing Assays” by Brad Larson (Agilent Technologies).

For gel invasion assays, a total of 5 000 A549 cells were seeded in low attachment U-bottom 96-well plates (Neolab, Heidelberg, Germany) with 100 µl of medium and centrifuged briefly for 2 minutes at 1 000 rpm to establish initial cell-cell contact. The cells were then incubated for 72 h under standard conditions. After this incubation period, the spheroids were gently collected into 1.5 ml Eppendorf tubes and allowed to settle on the tube’s bottom for 5 min on ice. Subsequently, a collagen G matrix gel was prepared, as described previously ^37^. Briefly, solution A was created by combining 1 M HEPES buffer (pH= 7-7.5) and 0.7 M NaOH in equal proportions. Solution A was mixed with 20% FBS in 10× PBS (pH = 7.4) in a 1:1 ratio to create solution B (pH = 7.90–8.05). For the final gelation step, collagen G and solution B were mixed in a 4:1 ratio (v:v). To create a collagen G+Matrigel matrix, a mixture of collagen G matrix gel and Matrigel was prepared in a 1:1 ratio (v:v). The spheroids were transferred into the matrix mixture and placed in an incubator to solidify over a period of 5 h. After solidification, the spheroids were imaged using a live cell imaging microscope for 72 h.

### Western Blot analysis

Denatured protein extracts were generated with the RIPA lysis buffer (50 mM Tris/HCl pH 7.5, 50 mM NaCl, 1% IGEPAL, 0.5% Sodium deoxycholate, 0.1% SDS). Briefly, frozen cell pellets were dissolved in RIPA buffer supplemented with protease inhibitor (Roche Diagnostics, Mannheim, Germany) and phosphatase inhibitor (Roche Diagnostics). The cell lysis process was carried out on ice and lasted for 20 minutes, with periodic vortexing during this period. Subsequently, the lysates were centrifuged at 15 000 rpm and 4°C for 20 minutes to remove cellular debris. Protein concentrations were calculated with BCA assay according to manufacturer’s recommendation (Pierce BCA Protein Assay Kit, Thermo Fisher Scientific) and 15 µg protein extracts were diluted with Milli-Q® water and mixed with 6x Laemmli buffer to a final concentration of 1x. Samples were denatured at 95 °C for 15 min. Polyacrylamide gels were prepared and cast using Bio-Rad equipment (Bio-Rad, Munich, Germany). Denatured protein extracts were loaded on the gel and electrophoresis was performed at 90 - 120 V.

Proteins were transferred to a methanol-activated PVDF membrane at 250 mA for 90 min at 4°C. Membranes were blocked with Roti®-Block (1 h) or EveryBlot (10 min). Primary antibodies were incubated overnight at 4°C, followed by three 15-minute PBST washes. HRP-conjugated secondary antibodies were applied for 1 hour at room temperature. After two PBST washes, chemiluminescence was detected using Luminata™ or SuperSignal reagents with iBright CL1500 (Thermo Fisher Scientific) or ChemiDoc (Bio-rad) systems. Densitometric analysis was conducted using ImageLab software.

**Table 1:**
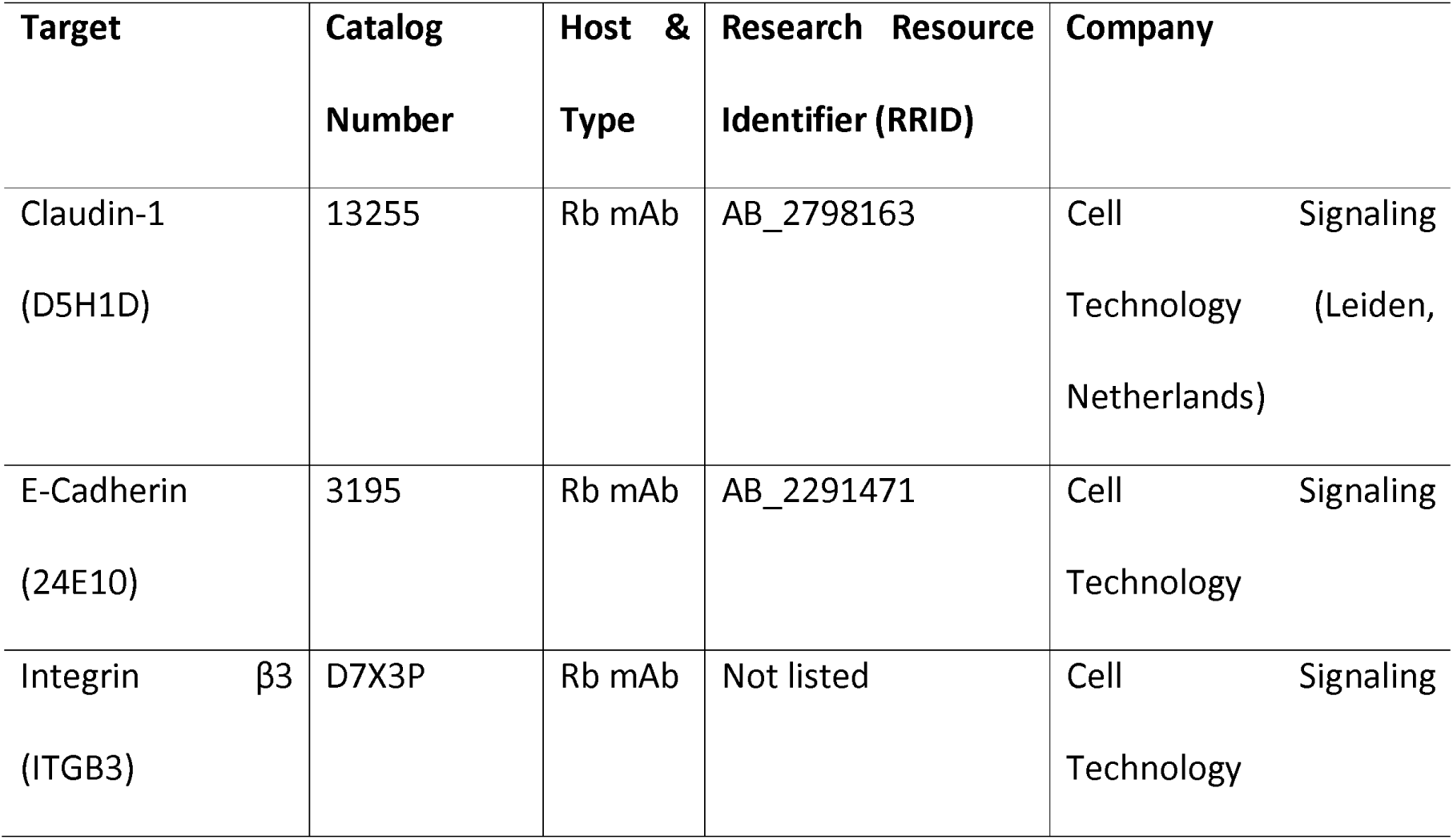
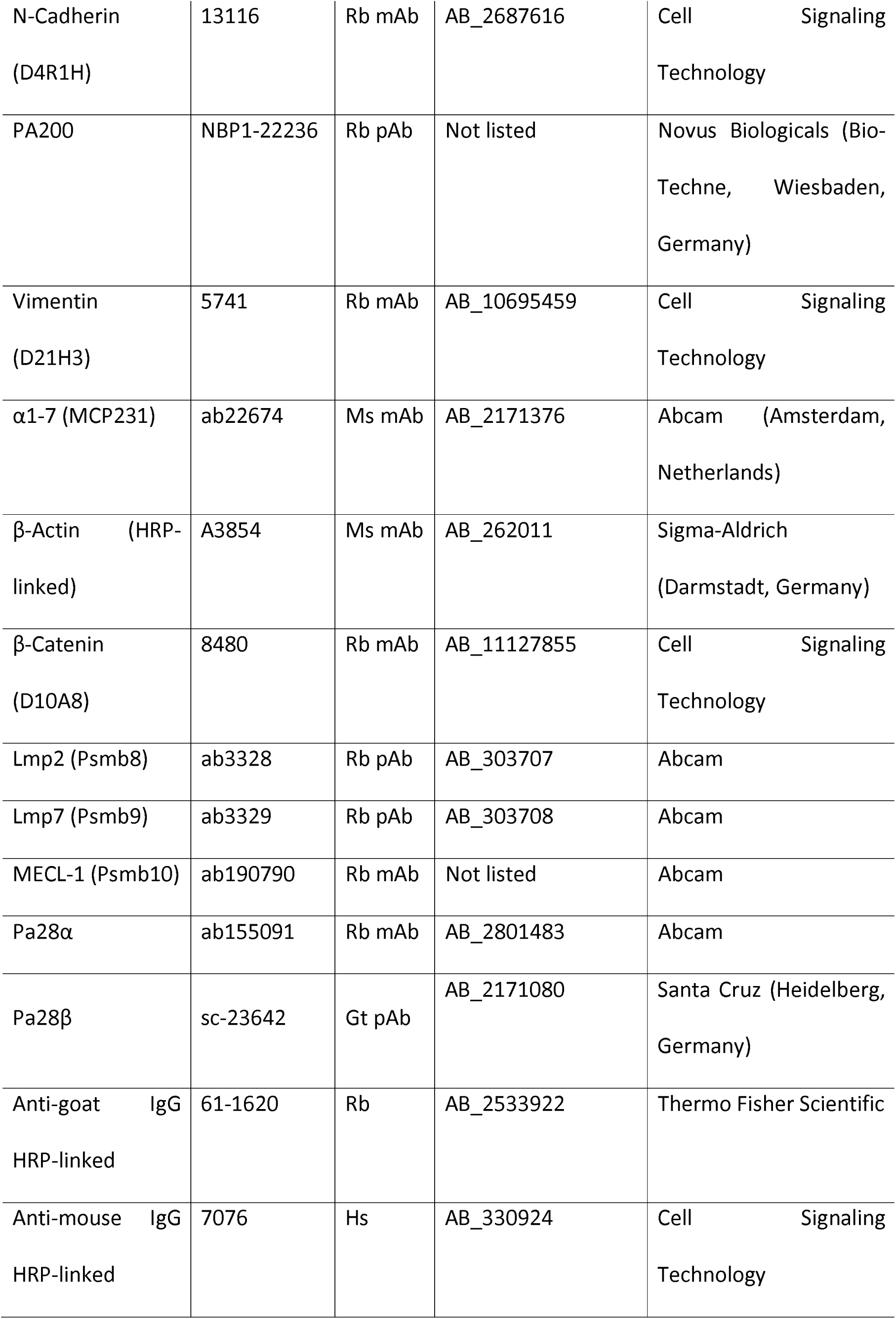
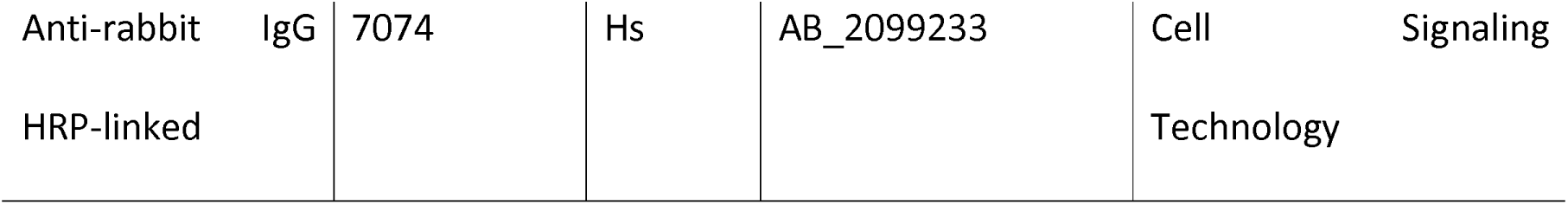
List of antibodies (Rb: Rabbit, Ms: Mouse, Gt: Goat, mAb: Monoclonal, pAb: Polyclonal)

### PA200 interactome analysis

For immunoprecipitation of PA200, cells were lysed in TSDG buffer (10 mM Tris pH 7.0, 10 mM NaCl, 1.1 mM MgCl_2_, 0.1 mM EDTA, 1 mM DTT, 1mM NaN_3_, 10 % (v/v) Glycerol) under native conditions as described previously to preserve physiological protein-protein interactions ^10,20^. Pre-clearing of the lysates was performed using magnetic Dynabeads coated with Protein A for antibodies raised in rabbits (Thermo Fisher Scientific) on a magnetic rack. 20 μL of Dynabeads were transferred to protein LoBind tubes (Eppendorf) and washed twice with 500 μL phosphate buffer pH 7.4. 200 μg protein lysate (in TSDG buffer + 0.2 % IGEPAL) was incubated with the Dynabeads for 20 minutes at room temperature on a rotator (20 rpm) in a pre-clearing step which allows the saturation of the unspecific binding sites of the Dynabeads with proteins from the cell extracts and thereby reduces unspecific background binding. In the meantime, 30 μl Dynabeads were washed twice with 100 μL phosphate buffer pH 7.4. Beads were resuspended in 50 μL phosphate buffer pH 7.4 and incubated with 3 μL of the anti-PA200 antibody (NBP1-22236, Novus) for 15 minutes at 1250 rpm and room temperature. Afterwards, beads were washed twice with 100 μL TSDG buffer + 0.2 % IGEPAL. 50 μl of the TSDG buffer + 0.2 % IGEPAL was used to resuspend the beads-antibody complexes. Precleared 200 μg TSDG protein lysates (separated from the beads) were added onto the beads bound to the antibody to a total volume of 250 μl. Samples were incubated with a rotator (20 rpm) for 2 hours at 4 °C. 25 μL per sample of the total mixture was used as input. 25 μl supernatant was collected once the beads had been separated on the magnetic rack. Input and supernatant samples (25 μl) were mixed with 6x loading buffer (5 μl) and heated at 95 °C for 10 min. Beads were washed five times (pre-cleared) with 500 μL TSDG buffer supplemented with 0.2 % IGEPAL and one final time with TSDG buffer. Co-immunoprecipitated proteins were eluted in 25 μL 1x Laemmli buffer and further denatured for 10 min at 95 °C. Eluted proteins were further analyzed by Western blotting together with the input and supernatant samples or sent to the Research Unit Protein Science (HMGU) for mass spectrometry analysis.

### In-gel proteasome activity assay

Native gel electrophoresis was performed to analyze intact and active proteasome complexes, following previously optimized and detailed protocol ^20^. Briefly, frozen cell pellets were thawed and lysed in OK lysis buffer (50 mM Tris/HCl pH 7.5, 2 mM DTT, 5 mM MgCl2, 10% glycerol, 2 mM ATP, 0.05% digitonin) supplemented with protease and phosphatase inhibitors (Roche Diagnostics). Cells were lysed on ice for 20 min with intermittent vortexing, followed by centrifugation at 15 000 rpm for 20 min at 4°C. Protein concentration was determined by BCA assay, and 15 µg of protein was diluted with water, mixed with 5x native gel loading buffer, and loaded onto 3-8% gradient NuPAGE Novex Tris-acetate gels (Themo Fisher Scientific). Electrophoresis was performed at 150 V for 4 h at 4°C in native gel running buffer. Following electrophoresis, an in-gel proteasome activity assay was conducted to assess chymotrypsin-like (CT-L) activity using an activity buffer (50 mM Tris, 1 mM ATP, 10 mM MgCl_2_, 1 mM DTT, and 0.05 mM Suc-LLVY-AMC). Fluorescence was detected using the ChemiDoc XRS+ system (Bio-Rad). For immunoblotting, gels were incubated in solubilization buffer (2% SDS, 66 mM Na_2_CO_3_, 1.5% β-mEtOH) for 15 min before protein transfer.

### Total proteasome activity assay

The activity of the 20S proteasome’s active sites—chymotrypsin-like (CT-L), caspase-like (C-L), and trypsin-like (T-L)—was evaluated using the Proteasome-Glo™ Assays (Promega, Fitchburg, Wisconsin, USA), following the manufacturer’s instructions. In brief, 1 µg of protein from OK lysates was mixed with OK lysis buffer (see above) to ensure equal volumes, and the final volume was adjusted to 20 µL with water. These dilutions were transferred to white flat-bottom 96-well plates and combined with 20 µL of the specific substrates. Luminescence was measured at 2-min intervals over 45 min using a Tristar LB 941 plate reader (Berthold Technologies, Bad Wilsbad, Germany). Each sample was analyzed in quadruplicate, and the plateau values of the luminescent signal were used for quantifying proteasome activity for each specific substrate.

### RNA isolation

For RT-qPCR, cellular RNA was isolated using the Roti®-Quick Kit (Roth, Karlsruhe, Germany) via phenol-chloroform extraction according to manufacturer’s instructions. Briefly, cells were lysed in Quick 1 solution, followed by the addition of Quick 2 solution. Samples were incubated on ice for 10 minutes, then centrifuged at 10 000 rpm and 4°C for 15 min to separate phases. The upper aqueous phase was carefully collected and transferred to a new tube and Quick 3 solution was added. The samples were incubated either at -80°C for 40 min or at -20°C overnight. RNA was pelleted by centrifugation at 13 000 rpm and 4°C for 20 min, followed by washing twice with 70% ethanol. The RNA pellet was air-dried on ice and reconstituted in 25-40 µL of nuclease-free water For bulk RNA sequencing, RNA was isolated using an RNA isolation kit (VWR, Darmstadt, Germany) according to the provided protocol. Cell pellets were lysed in TRK buffer with 2% β-mEtOH and homogenized using a peqGOLD RNA Homogenizer Column (VWR). Following centrifugation at ≥12 000 x g for 2 min, the flow-through was mixed with ethanol (70%) and vortexed. The mixture was then passed through a peqGOLD RNA Mini Column via centrifugation at 10 000 x g for 1 min. The column was washed with RNA Wash Buffer I and 80% ethanol, followed by centrifugation steps to remove residual ethanol. RNA was eluted by nuclease-free water and concentration was determined using the NanoDrop 1000 (Thermo Fisher Scientific).

### Reverse transcription of RNA and RT-qPCR

For the reverse transcription process, RNA ranging from 0.25 to 1 µg was mixed with 9 µL of nuclease-free water, followed by the addition of 2 µL of 250 µM Random Hexamers (Thermo Fisher Scientific). After allowing the mixture to incubate at 70 °C for 10 min, the samples were placed on ice. Subsequently, 9 µL of a reverse transcription master mix was introduced, with final concentrations of 1x First Strand Buffer, 10 mM DTT, 0.5 mM dNTPs, 1 U/µL RNAse Inhibitor, and 10 U/µL M-MLV transcriptase. The reverse transcription procedure involved annealing for 5 min at 25°C, followed by elongation for 60 min at 37°C, which was executed using a Mastercycler Nexus (Eppendorf, Hamburg, Germany). The cDNA was subjected to digestion with 1 U of DNase at 37°C for 15 min, followed by heat inactivation at 75°C for 10 min. The resultant cDNA was then diluted at a 1:5 ratio with nuclease-free water (Ambion, Thermo Fisher Scientific).

Quantitative real-time RT-PCR analysis was carried out using the SYBR Green LC480 system from Roche Diagnostics. A mixture comprising 2.5 µL of both forward and reverse primer dilutions (0.5 µM each) and 5 µL of LC480 SYBR Green I Master mix (Roche Diagnostics) was prepared per well in a 96-well plate format. Additionally, 2.5 µL of cDNA was added to each well. Each sample was measured in duplicate, and the plates were centrifuged for 2 minutes at 1 000 rpm prior to initiating the measurements using the standard program outlined in Table 4.7 on the Light Cycler 480 II (Roche Diagnostics). The gene expression levels of the various samples were normalized to the housekeeping gene ribosomal protein L19 (RPL19). The relative gene expression was determined using the ΔΔCT method, with the specificity of the primers verified by analyzing the melting curve.

**Table 2:**
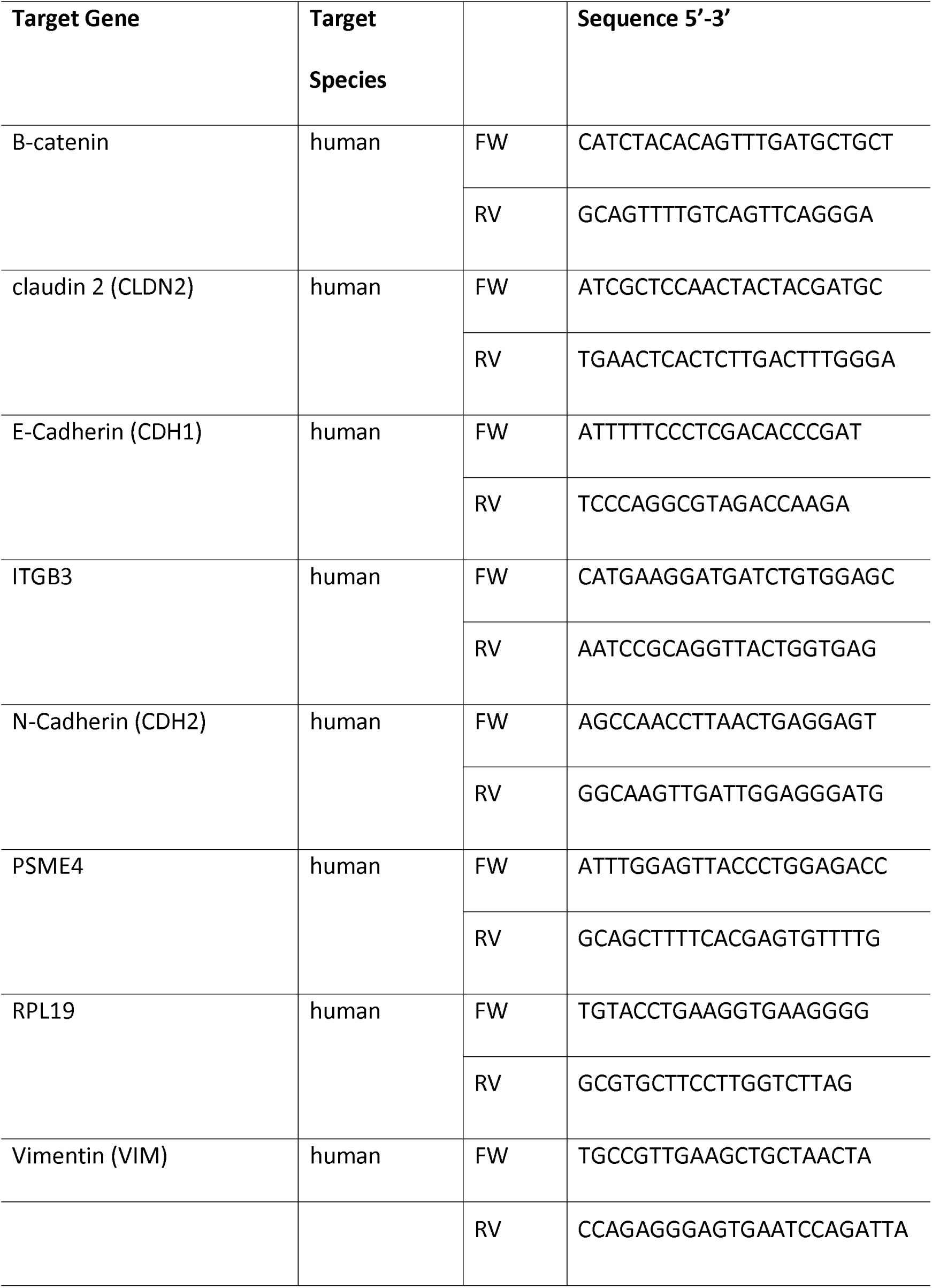
List of primers (FW: Forward primer, RV: Reverse Primer)

### Bulk mRNA sequencing and analysis

A total of 1.5 million A549 cells and 1 million H1299 cells were cultured in 15 cm dishes and incubated for 48 hours. Total RNA was isolated utilizing the Total RNA kit (Peqlab, VWR). Subsequently, the samples were dispatched to the Core Facility Next-Generation Sequencing at the Helmholtz Center Munich. The RNA integrity number (RIN) was assessed using the Agilent 2100 Bioanalyzer system, and concentration was quantified using the Qubit 2.0 RNA BR assay (Invitrogen). RNAs with RIN values exceeding 6 were chosen for total RNA sequencing (ribo-depleted), a method that effectively detects essential coding and non-coding transcripts by removing highly abundant rRNA species.

Library preparation for both human and mouse samples involved using the TruSeq stranded total-RNA Library Preparation kit (Illumina) with 1 μg of RNA, following the kit’s protocol. Following a final quality control step, the libraries were subjected to paired-end sequencing (2x150 bases) on the Hiseq 4000 sequencer (Illumina), targeting a depth of at least 40 million paired reads per sample. Subsequent to sequencing, initial processing (demultiplexing) of raw data to fastq files was performed using the bcl2fastq v2.20 program from Illumina.

For RNA sequencing file alignment, the STAR tool was employed with the GRCh38/hg38 version. To estimate transcript abundances, the FeatureCounts function within the Subread2 package was utilized in the Unix environment. Initially, the gene-code ReleaseM23 (GRCm38.p6) served as the gene annotation. Subsequently, data underwent filtering, normalization, and differential gene expression analysis using relevant packages within the R programming environment.

limma: (https://bioconductor.org/packages/release/bioc/html/limma.html)

edgeR: (https://bioconductor.org/packages/release/bioc/html/edgeR.html)

### Animal experiments

The experimental procedures involved NOD/SCID mice purchased from Charles River (Freiburg, Germany) (RRID:IMSR_CRL:394) and experiments were performed according to local regulations in Bavaria, Germany (TVA approval number ROB-55.2-2532.Vet_02-20-197). Male and female animals of 6-8 weeks of age were used in the experiments, the sample size was determined beforehand through a G*power analysis. We used G*Power v3.1.9.7. (http://www.gpower.hhu.de/; accessed on 14.10.2020, RRID:SCR_013726) and calculated α = 0.054 and β = 0.217 as acceptable sizes. According to this calculation and to demonstrate an effect size of f > 0.05 with α < 0.05 and β < 0.20, we needed at least a sample size of n = 10 per group. We therefore used 12 mice per group, taking into account 20% reserve animals. Experimental procedures were randomized across different cages. The animals were anaesthetized before being injected with 3x10^6^ PA200 WT or PA200 KO cells in 150 µl PBS, and their body weight and tumor size were measured regularly. Mice had to be sacrificed once their tumor burden reached the maximal size, the primary tumor had necrosis or 10 weeks after the cell injection. Once the endpoint conditions were met, the mice were euthanized using lethal anesthesia for organ harvesting and further examinations.

### Immunohistochemical analysis of mouse and human tissues

Excised mice tissue specimens were fixed in a 4% (w/v) solution of formalin, followed by paraffin embedding and sectioning into slices measuring 3 µm thick. These sections were subsequently stained using hematoxylin and eosin (HE) staining. The staining procedure was carried out using the HistoCore SPECTRA ST automated slide stainer (Leica, Wetzlar, Germany) with ready-made staining reagents (Histocore Spectra H&E Stain System S1, Leica) following the instructions provided by the manufacturer. Immunohistochemical staining was performed following standardized protocols on a Discovery Ultra automated stainer (Ventana Medical Systems, Tucson, Arizona, USA). Polyclonal rat anti-Ki67 (1:1 000, Abcam) or anti-PSME4 (sc-135512, Santa Cruz, RRID:AB_2171430) was used as a primary antibody. Signal detection was achieved using the Discovery® DAB Map Kit (Ventana Medical Systems, Tucson, AZ). The stained tissue sections were then digitally scanned using an AxioScan.Z1 digital slide scanner (Zeiss) equipped with a 20x magnification objective.

Human lung tissue obtained from patients surgically treated for lung cancer was provided by the Asklepios Biobank for Lung Disease, Gauting, Germany. Samples were approved with ethical consent obtained from the ethics committee of Ludwig-Maximilians University Munich, in accordance with national and international guidelines (project number 333-10). Paraffin embedded human tissues were cut in 3 μm thick sections using the Hyrax M55 microtome (Zeiss). Tissue sections were incubated for one hour at 60 °C in order to melt paraffin, deparaffinized by incubating two times in xylene for 5 min and rehydrated in a descending alcohol series (100 %, 90 %, 80 % and 70 % (v/v)) for 1 min. To block endogenous protease activity and to permeabilize sections for nuclear staining they were incubated in a methanol/hydrogen peroxide (80 %/1.8 % (v/v)) solution for 20 min. Tissue sections were rinsed in Milli-Q® water and heat-induced antigen retrieval was performed in citrate buffer pH 6 using a decloaking chamber (Biocare Medical, Zytomed Berlin, Germany). After washing with TBST unspecific binding sites were blocked for 30 min with Rodent Block M (Biocare Medical). The slides were washed again in TBST and incubated with anti-PA200 antibody (sc-135512, Santa Cruz, RRID:AB_2171430) diluted in Antibody Diluent (DAKO) for 1 h at RT. After extensive washing in TBST, sections were incubated with MACH 2 Rabbit AP-Polymer (Biocare Medical) for 30 min at RT. Sections were rinsed again in TBST and incubated in Vulcan Fast Red AP substrate solution (Biocare Medical) for 10 min. Tissue sections were washed in TBST and MilliQ® water and hematoxylin counterstaining (Roth) was performed to visualize nuclei. After repeated washing in TBST, sections were dehydrated in ethanol and xylene and mounted using Entellan mounting medium (Merck Millipore, Darmstadt, Germany). Slides were imaged using the MIRAX scanning system (Zeiss). Stainings were analyzed by an expert clinical pathologist blinded to the sample identity. Semiquantitative scores for PA200 expression were obtained by defining the percentage of PA200 positively stained tumor areas multiplied by the intensity of staining as graded between 1 (weak) to 3 (strong). Scores were dichotomized into high (>80) and low (<80) expressing tumors with the score 80 representing the median of all samples and PA200 expression scores were correlated with survival of the patients for this cohort.

### Statistical analysis

The specific statistical analyses for each panel are provided in the corresponding figure legends. Statistical significance was denoted in the figures as follows: * for p < 0.05, ** for p < 0.01, *** for p < 0.001, or **** for p < 0.0001. The figures display data as mean ± SEM. All statistical analyses were performed using GraphPad Prism software (version 10, RRID:SCR_002798).

## ACKNOWLEDGEMENT

This work was supported by the Leibniz Association granted to SM. We are grateful for the technical support of Core Facility Genomics at Helmholtz Zentrum München. We also like to thank Elvira Stacher-Priehse from the Askepios Klinik Gauting for her analysis of the histological stainings.

## AUTHOR CONTRIBUTION STATEMENT

ASY, FK, LZ, GGG, SJB, VW, TM performed experiments; ASY, LZ, GGG, SJB, TG, GTS, VW and NR analyzed the data; ASY, SM wrote and revised the manuscript; SM developed and coordinated the project.

## ETHICS STATEMENT

Animal experiments were performed according to local regulations in Bavaria, Germany (TVA approval number ROB-55.2-2532.Vet_02-20-197). The maximal tumor size permitted by the ethics committee was 20 mm. Once the tumor size approached this volume, mice were sacrificed. Human lung tissue obtained from patients surgically treated for lung cancer was provided by the Asklepios Biobank for Lung Disease, Gauting, Germany. Samples were approved with ethical consent obtained from the ethics committee of Ludwig-Maximilians University Munich, in accordance with national and international guidelines such as the Declaration of Helsinki (project number 333-10).

## DATA AVAILABILITY

The data generated in this study have been made publicly available in Gene Expression Omnibus under GSE288117 (GEO, RRID:SCR_005012).

