## Supplemental Information for "PA200 differentially regulates the proteasome and inhibits migration of cancer cells"

This supplemental information contains additional figures and their legends to complement data sets that are not included in the main manuscript.

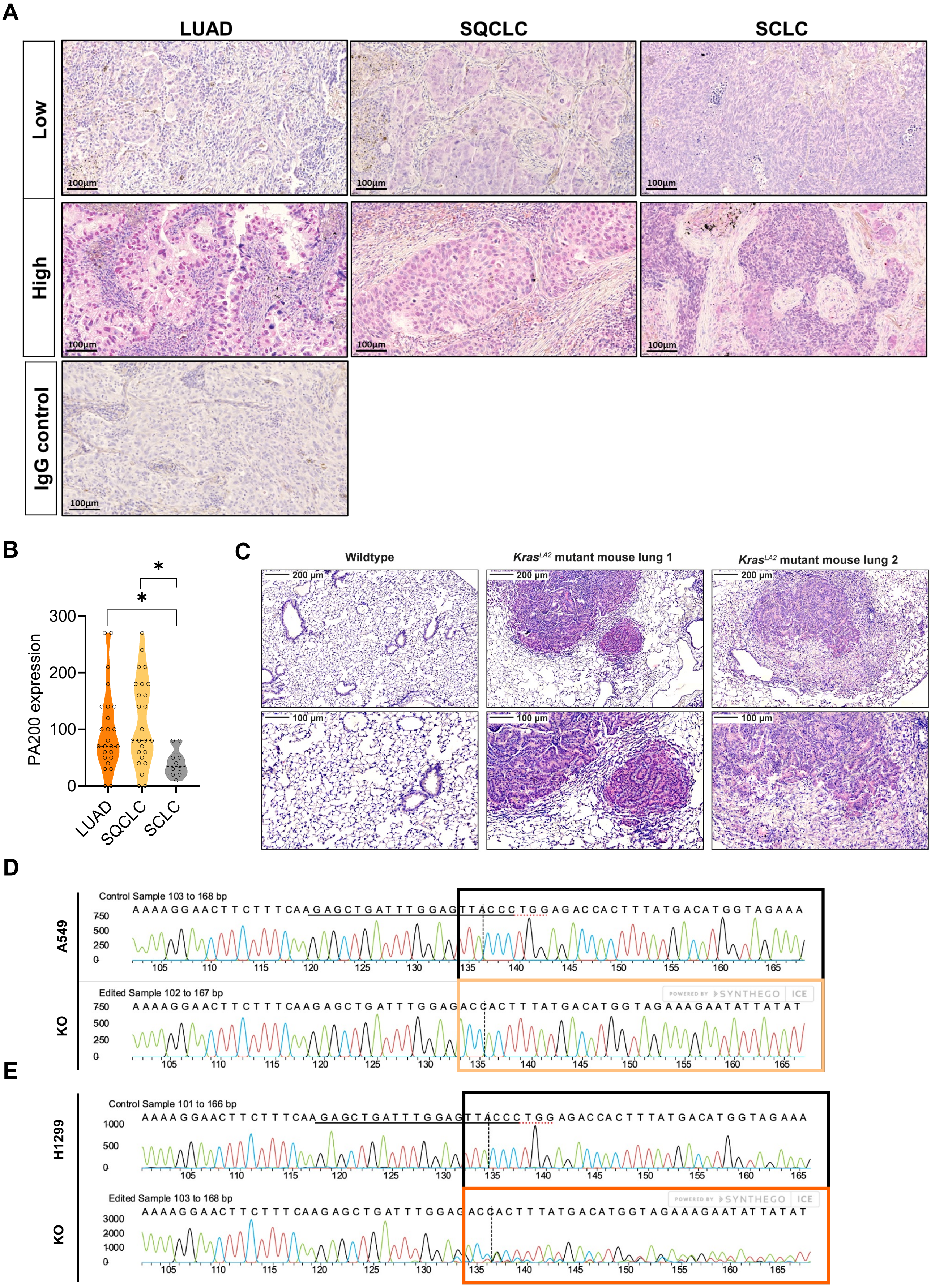


**Supplementary Figure 1: PA200 is upregulated in NSCLC tumors. A)** Representative images of tumor sections in LUAD (Lung adenocarcinoma), SQCLC (squamous cell lung cancer), and SCLC (small cell lung cancer) and IgG control in LUAD tissue. **B)** Immunohistochemical analysis of tumor sections (24 LUAD, 14 SQCLC, 9 SCLC), semiquantitatively graded by a blinded pathologist (Kruskal Wallis, Dunn's multiple comparisons test). **C)** Expression of PA200 (pink) was determined in wildtype and KrasLA2 mutant mouse lungs by immunohistochemistry. Nuclei were counterstained with hematoxylin (blue). Sanger sequencing results were aligned with a non-edited DNA template from **D)** A549 and **E)** H1299. The guiding target sequence is underlined in the WT sequence and vertical dotted black lines indicate the anticipated cut place.

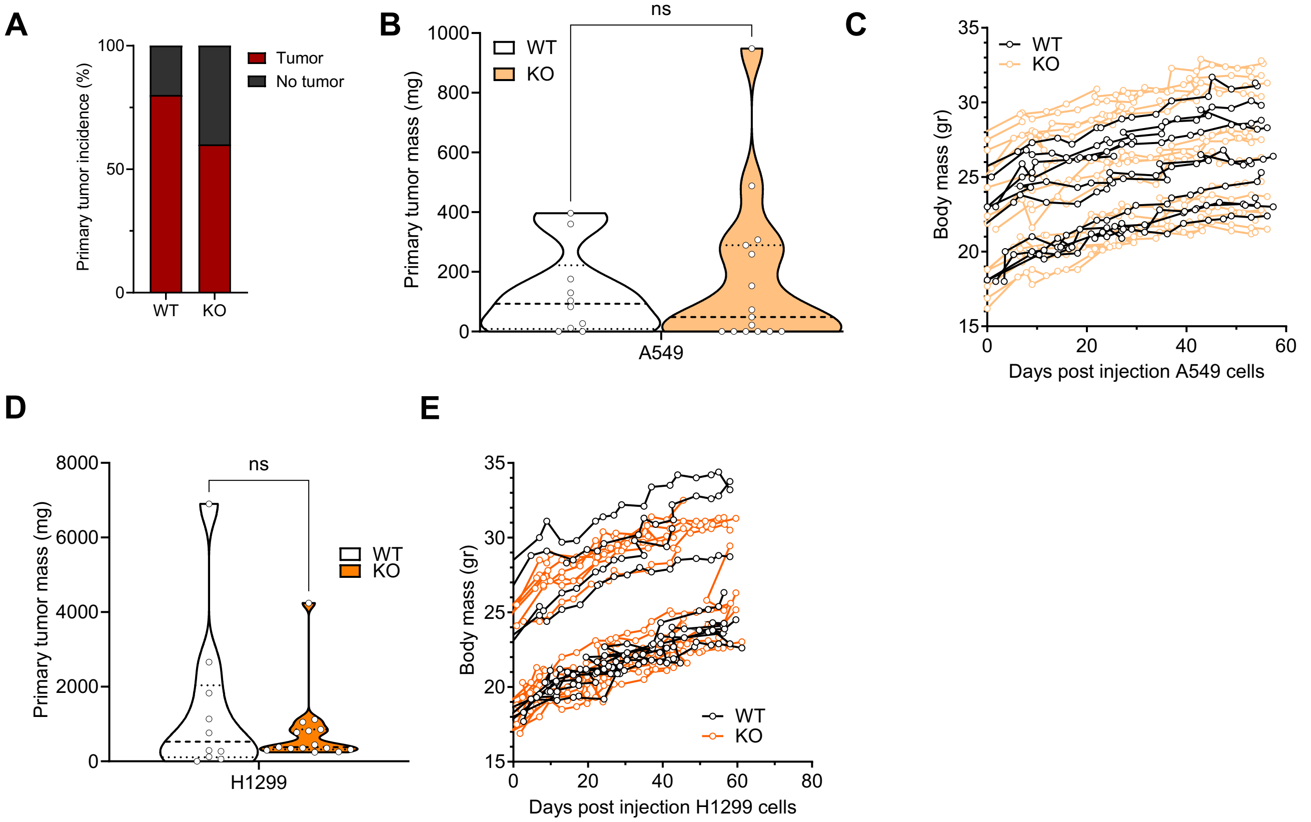


**Supplementary Figure 2: Deletion of PA200 in lung cancer cell lines does not consistently alter tumor or body mass. A)** Primary tumor incidence for A549 cells. **B,D)** Primary tumor mass (mg) was weighed after dissection, and **C, E**) the body mass of the mice was tracked over the course of the experiment for **B-C)** A549 and **D-E)** H1299 injected mice.

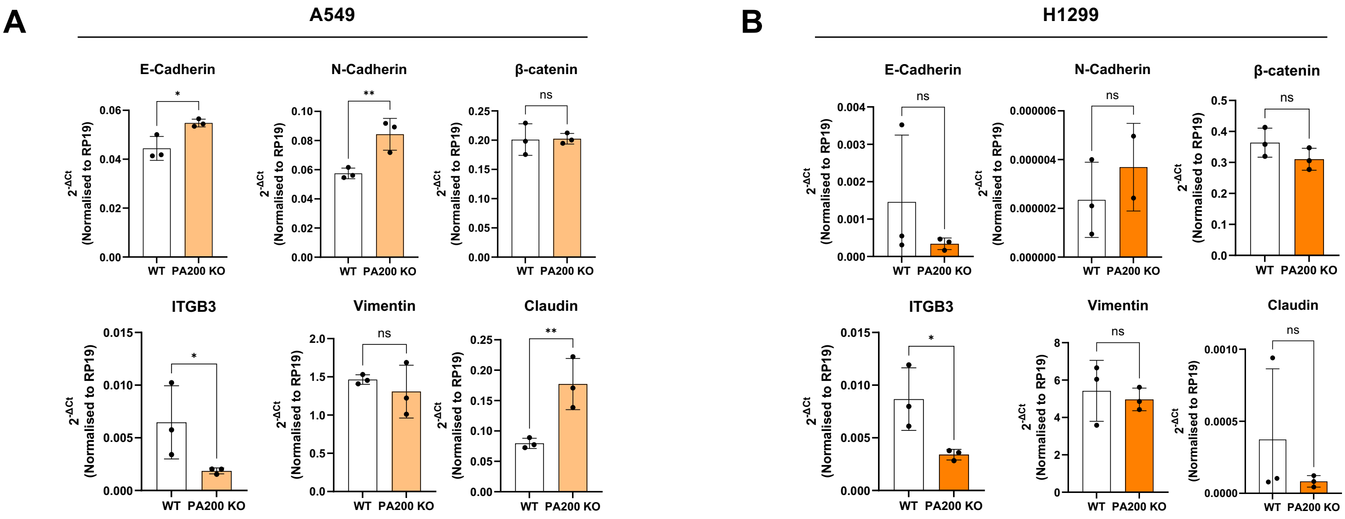


**Supplementary Figure 3: Deletion of PA200 impacts mRNA levels of EMT markers.** mRNA levels of epithelial and mesenchymal markers were quantified (right panel, n=3, different passages, 2-way ANOVA) in **A)** A549 and **B)** H1299 WT and KO pooled cell clones and RPL19 used as housekeeping gene. For quantification, relative expression levels were normalized to the respective WT control of each cell line. *p < 0.05, **p < 0.01, ***p < 0.001, ****p < 0.0001.

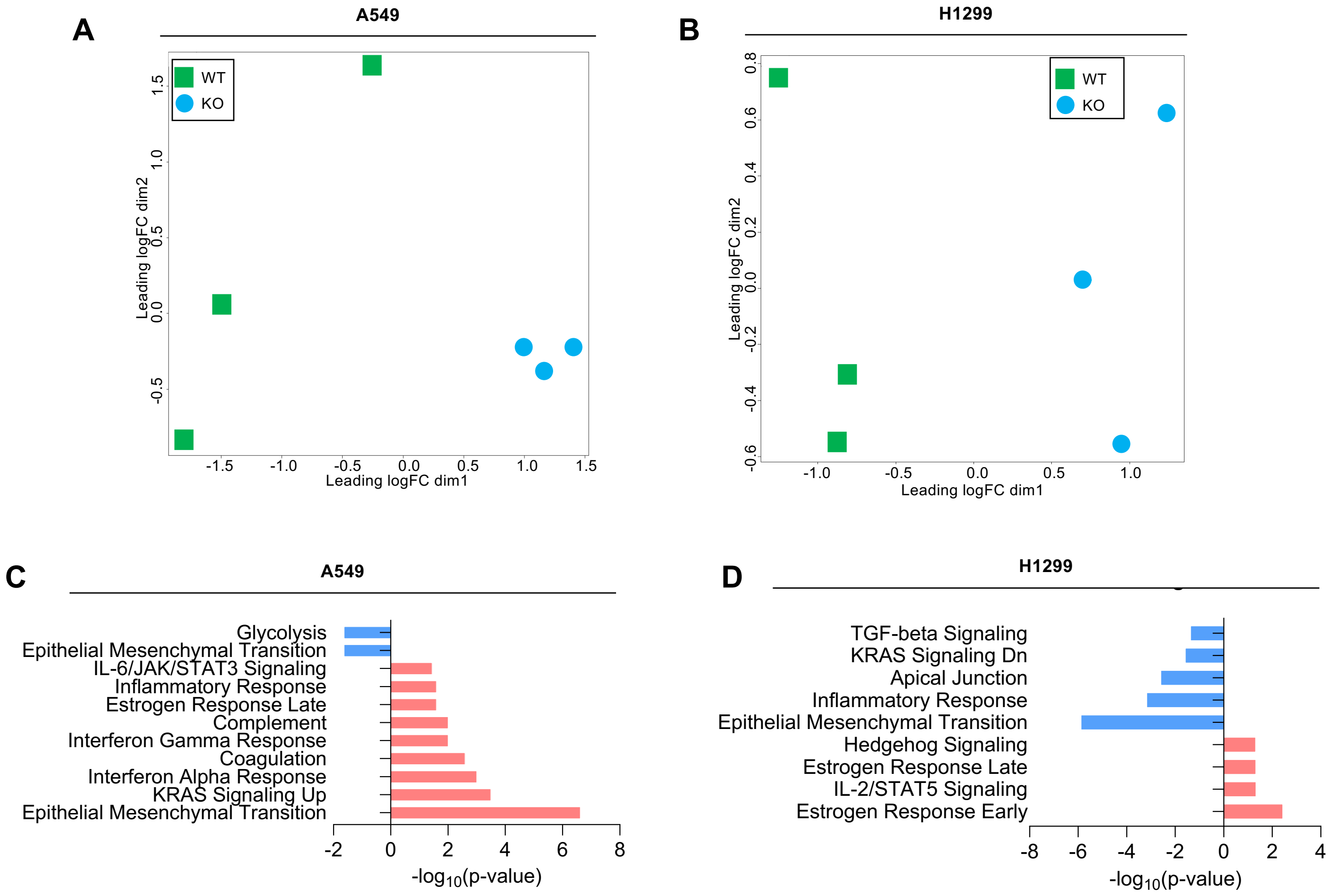


**Supplementary Figure 4: Transcriptome analysis of A549 and H1299 WT and KO clones.** Principal component analysis (PCA) of **A)** A549 and **B)** H1299 cell clones. Significantly up- or down-regulated genes were used for further analysis, and only the top 10 results of MSigDb are shown in the graphs for **C)** A549 and **D)** H1299.

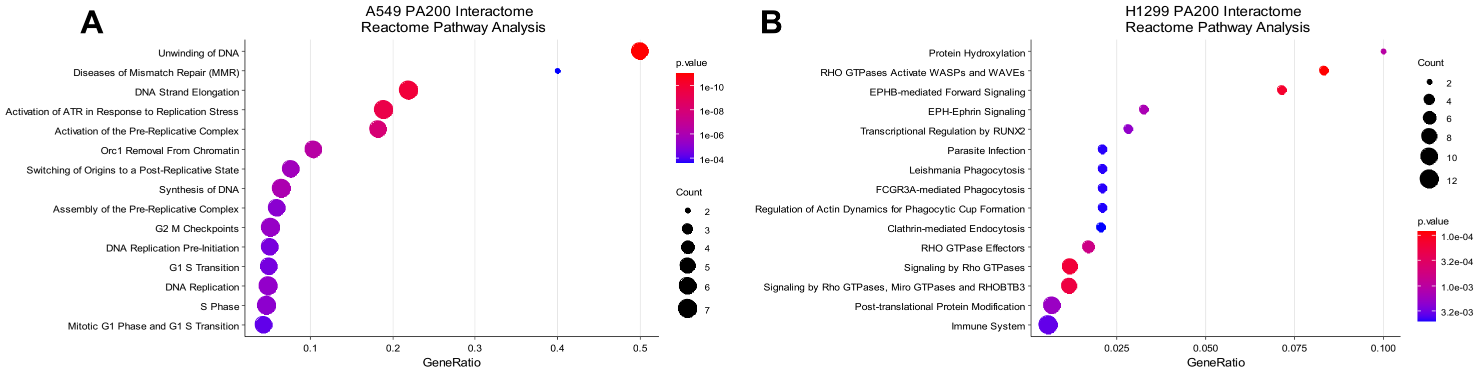


**Supplementary Figure 5: Interacting proteins of PA200 in A549 and H1299 cells with diverse functions.** Bubble charts of Reactome Pathways analysis of PA200 interactors in **A)** A549 and **B)** H1299 cells.
